# nf-core/genomeqc: a best-practice pipeline for comparing genome and assembly quality

**DOI:** 10.64898/2026.08.20.745971

**Authors:** Christopher D.R. Wyatt, Fernando Duarte Frutos, Stephen D. Turner, Alex Cerqueira De Araujo, Usman Rashid, Victoria Begley, Seirian Sumner

## Abstract

The rapid growth in publicly available genome assemblies has made selecting genomes suitable for downstream analyses increasingly challenging. Differences in assembly and annotation quality can influence gene completeness, duplication rates, contiguity, repeat representation, and other characteristics. Assessing genome quality therefore requires integrating multiple complementary quality metrics that are often generated by independent tools. Here, we present **nf-core/genomeqc**, a workflow for assessing and comparing genome assemblies. The pipeline accepts RefSeq/GenBank accessions for automatic genome and annotation retrieval, or local genome (FASTA) and annotation (GFF3/GTF) files. It integrates complementary analyses of assembly contiguity, gene completeness, annotation quality, repeat content and other quality metrics using tools such as BUSCO, QUAST, Merqury, and AGAT, before combining the results on a phylogenetic tree for visualisation and comparison across species. GenomeQC is implemented in Nextflow within the nf-core framework, providing an accessible, reproducible, scalable and community-driven workflow for genome quality assessment.

## Introduction

Not all genome assemblies and annotations are created equal. Genome quality can vary dramatically between species and studies depending on sequencing technology (short-read vs long-read vs hybrid)(1), assembly algorithms and parameterisation, annotation pipelines(2) and tools, and even the individual/tissue sequenced because DNA integrity(3), contamination and ploidy/heterozygosity differ between samples(4). Without accounting for these differences, we risk biasing our analyses and drawing incorrect conclusions from comparative genomic data(5). Therefore, assessing genome and annotation quality is a prerequisite for meaningful comparative genomic analyses.

### Assembly quality determines how accurately a genome is reconstructed

Assemblies vary in completeness, contiguity and sequence accuracy. Fragmented assemblies obscure synteny and gene order (2, 4); repetitive telomeric and centromeric regions are often collapsed or missing (4); and organellar (mtDNA) genomes are inconsistently captured across databases (6). The assembly algorithm itself shapes how repeats and segmental duplications are represented (2), while platform-specific sequencing errors introduce false variants and frameshifts(1) and heterozygosity can split alleles into duplicate loci, inflating apparent gene counts(7). Each of these is quantifiable through contiguity statistics, single-copy orthologue completeness, telomere content, k-mer-based accuracy and contamination screening, but are considerably more informative when interpreted alongside closely related genomes.

### Annotation quality represents a second, distinct source of variation

Even high-quality assemblies can produce markedly different annotations depending on the annotation pipeline, the completeness and tissue representation of RNA-seq evidence, and the reference databases used. Gene counts, pseudogene identification, exon–intron boundaries and isoform prediction all vary between annotation approaches (8–12). As a result, apparent differences in gene content between species may reflect annotation methodology rather than genuine biological differences.

Several tools are available for assessing genome quality, including cogeqc(11), Blobtoolkit(13), AssemblyQC(14) and HuffordLab/GenomeQC(15). These provide valuable metrics for evaluating assembly and annotation quality, including measures such as BUSCO completeness, assembly contiguity, scaffold statistics and GC content. However, these metrics are typically reported for individual assemblies, making it difficult to place genome quality into a broader comparative context. nf-core/genomeqc integrates these core analyses with additional assessments, including RepeatMasker-based repeat content analysis, gene overlap detection, BUSCO ideograms(16) and TIDK-based telomere detection, and presents all quality metrics within a shared phylogenetic framework, allowing assemblies to be interpreted relative to closely related species. Implemented within the nf-core community(17), the pipeline benefits from community-driven development, reproducibility and long-term maintenance. A comparison of feature overlap is given in **Supplementary Table S1**.

Here we present **nf-core/genomeqc** v1.0.0, a Nextflow pipeline for comprehensive comparative assessment of genome assemblies and annotations. Rather than reporting quality metrics in isolation, **genomeqc** integrates multiple complementary analyses and visualises them within a phylogenetic context, enabling users to evaluate genomes relative to closely related species.

## Materials and Methods

### Implementation and reproducibility

**nf-core/genomeqc** is written in Nextflow(18), following the standards of the nf-core(17) community. It is a modular pipeline built using nf-core modules where possible (many of which were built at the October 2024 nf-core hackathon). These are unit-tested and containerised so that each tool runs without local installation. The nf-core framework keeps the pipeline readable, standardised and open to community contribution. The code is hosted in the nf-core GitHub repository (https://github.com/nf-core/genomeqc/), enabling contributions and discussion of bugs, issues and feature requests, and tagged releases provide version tracking and reproducibility. Configuration profiles support a range of container engines and package managers, e.g. Docker, Singularity, Podman, Apptainer and Conda, as well as user-supplied institutional profiles for specific HPC or cloud environments (https://nf-co.re/configs/; usage: -profile <institution>).

Running the pipeline requires a container engine or conda, Nextflow v26.6.4 or later and Java v17–25. All individual programs run by the pipeline are cited in **Supplementary Table S1**. At runtime, every tool and version used in a given run is written to a software version yml file in the results/pipeline_info/ folder, allowing users to cite the exact tools applied to their data. A custom run is specified with an input comma-separated samplesheet (e.g. --input mysamplesheet.csv) (**Table 1**) listing either RefSeq/GenBank IDs or paths to input genomes (FASTA) and, optionally, annotations (GFF/GTF).

**Table 1.** Structure of the input samplesheet for nf-core/genomeqc. *A*: An example samplesheet. Values are comma-separated and a header line is always required. Each row must contain a unique identifier (species) and a genome, supplied either as an NCBI accession (ncbi) for automatic download or as a path to a local assembly (fasta). The remaining columns are optional: an annotation file (gff), long reads for Merqury assembly evaluation (fastq), and an NCBI taxid for foreign contamination screening (taxid). Unused fields are left empty. Columns unused by every sample may be omitted entirely; a samplesheet containing only species and ncbi, for instance, is valid. Working examples can be found at https://github.com/Eco-Flow/test-datasets/blob/genomeqc/samplesheet/. ***B***: Description of each column in the samplesheet.

| Column | Description |
| --- | --- |
| species | Species name or custom sample name. Spaces in sample names are automatically converted to underscores ( _ ). |
| ncbi | ncbi accession. Can be GenBank (starts with GCA ) or RefSeq (starts with GCF ). |
| fasta | Full path to the genome FASTA file. Can be compressed or uncompressed. |
| gff | Full path to the genome annotation GFF/GTF file. Can be compressed or uncompressed. |
| fastq | Full path to FASTQ file for long reads (e.g. PacBio or ONT). File has to be gzipped and have the extension ".fastq.gz" or ".fq.gz". |
| taxid | Species taxid for decontamination screening, must be a valid NCBI taxid (numeric string without spaces). |

### Test data

Test data is packaged with the pipeline, and can be run with ‘nextflow run main.nf -profile docker,test’ (for example), which pulls a tiny test dataset composed of four mycoplasma genomes from a samplesheet in the repository. To confirm correct operation of the pipeline, the small test dataset will run through all steps (except the optional read-based Merqury and Tiara steps, which require raw genome data). This ensures that the full run completes within minutes during continuous integration tests set up in Github actions, and we can ensure future changes do not break the pipeline, as nf-test(19) snapshots are kept within the repository. **Supplementary Table S2** contains the mycoplasma related species sample sheet used in the following figures.

## Results

### Pipeline overview

**nf-core/genomeqc** takes two or more genomes, supplied as local FASTA/GFF files or NCBI RefSeq/GenBank accessions (or a mixture) from a single CSV samplesheet (**Supplementary Table S2**), and runs a set of quality-assessment tools whose outputs are unified for direct comparison (**Figure 1**). Analyses fall into three tracks: genome-only metrics (blue in **Figure 1**), annotation-dependent metrics (green), and steps requiring both (red). The pipeline runs only the tracks supported by each input, skipping gene-level analyses where no annotation is provided. Some steps are optional, such as the contamination (FCS (20), Tiara(21)) and transposon characterisation (Repeatmasker(22) / Hite(23)). Genomic read-based (Merqury) steps are triggered by supplying raw reads from the genome construction. Results are collated into a MultiQC report (24), custom HTML and Excel summary tables, and an annotated phylogenetic tree (**Figure 3**) giving a comparative view across all genomes; the Supplementary Information describes each tool in detail.

**Figure 1.**
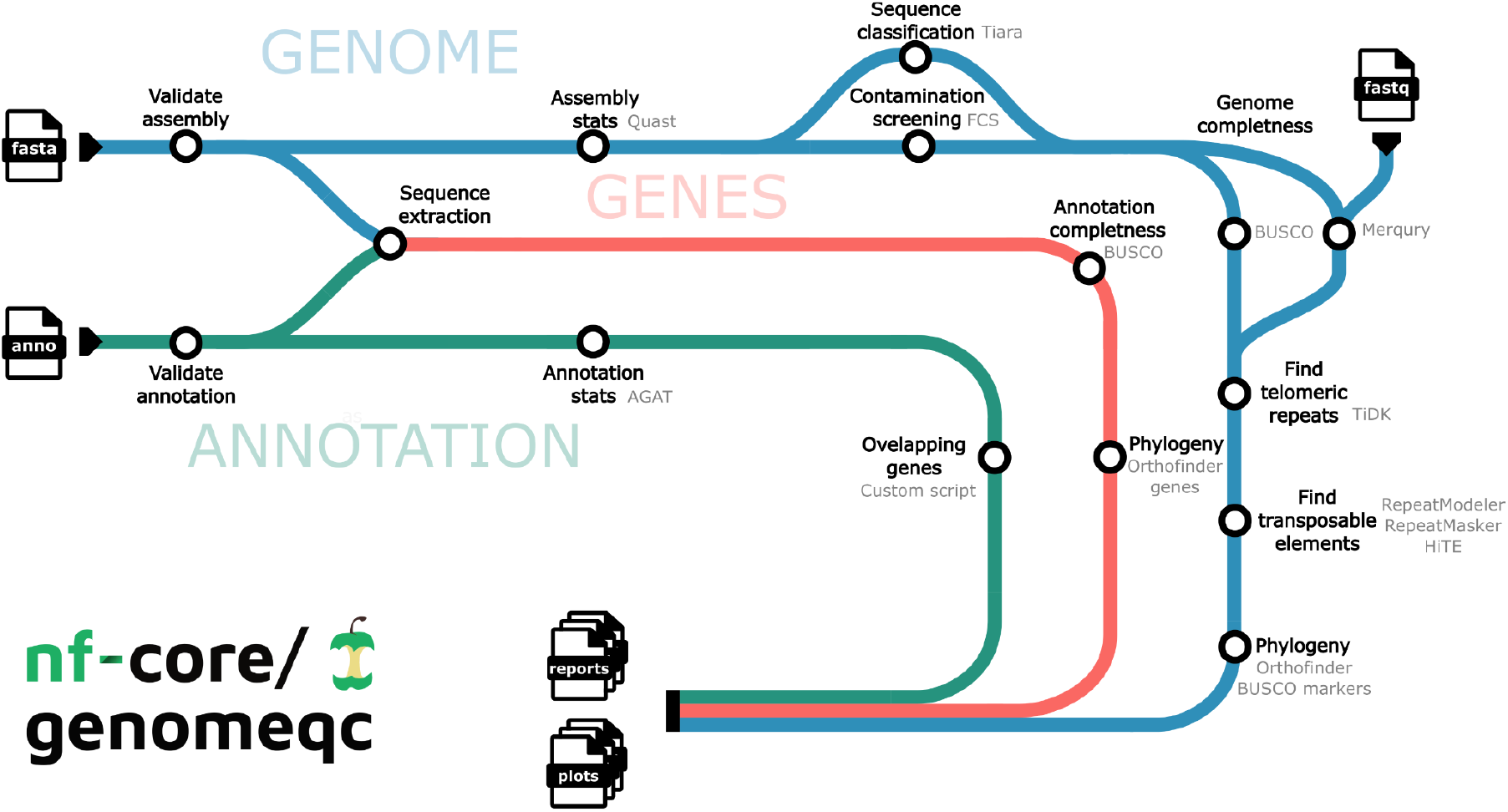
Pipeline overview. Tube map of nf-core/genomeqc (v1.0.0) steps. The pipeline is divided in three labelled sections (genome only tasks, annotation only tasks and tasks related to genes), with each node representing a specific tool. Input files are listed in page icon symbols with their respective input type labelled.

### Comparative phylogenetic report

To demonstrate the pipeline, we ran 15 Mycoplasma species (**Table S2**), spanning a range of assembly qualities, through the full workflow. The central output is a phylogeny that annotates every assembly with its quality metrics side by side, available in two forms: a circular layout (**Figure 2**) and a traditional, left-aligned rooted layout (**Figure 3**). When annotations are supplied, the tree is inferred from orthogroups identified by OrthoFinder(25) using the longest protein isoform per gene; with genomes alone, it is built from shared BUSCO single-copy markers(26). At least three assemblies are required to construct the tree. This comparative framing helps spot an outlier caused by assembly or annotation artefacts from genuine biological variation, a distinction that is hard to make from per-genome reports in isolation.

**Figure 2.**
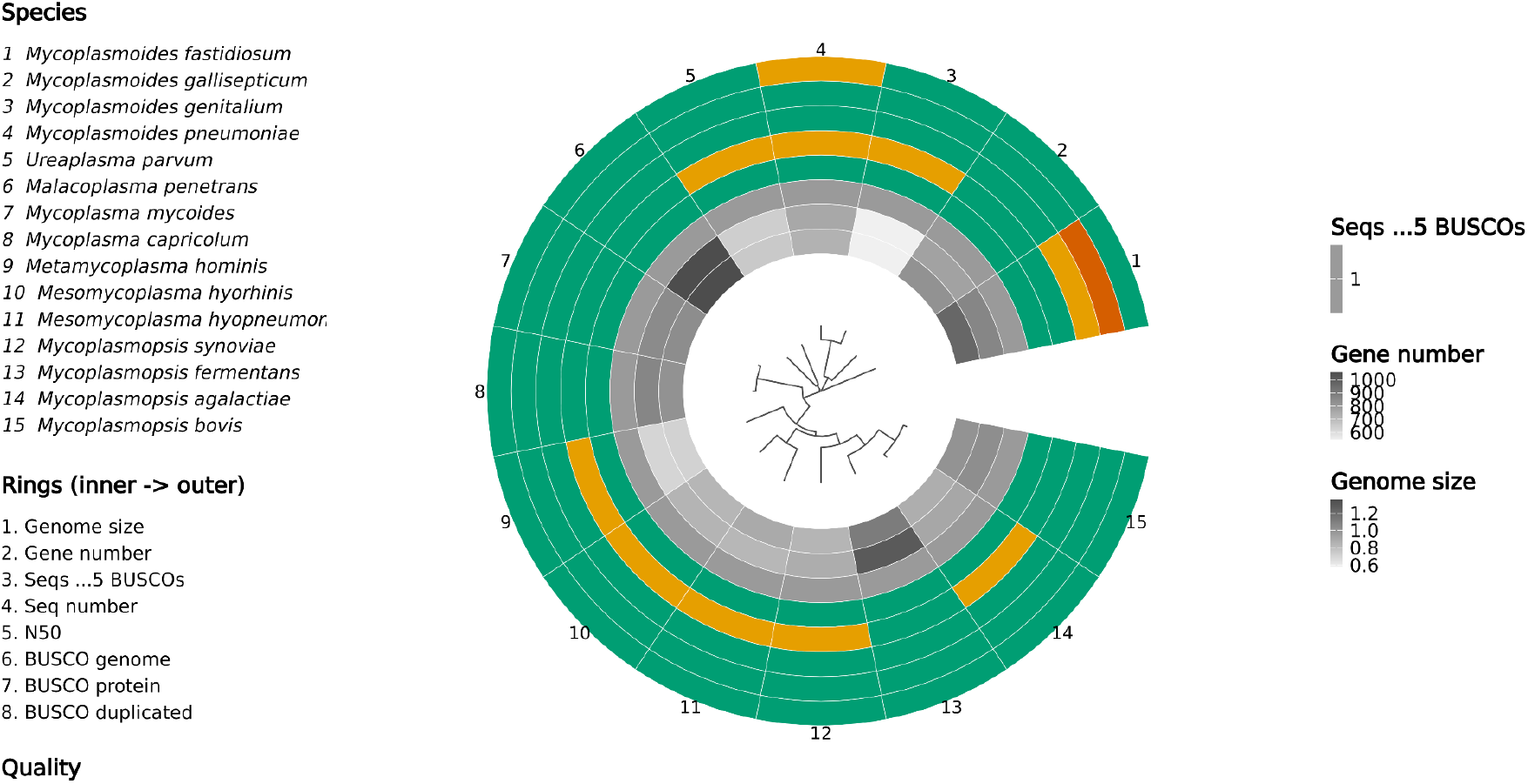
Circular phylogeny with QC rings. A phylogeny inferred from OrthoFinder orthogroups (or, genome-only, shared BUSCO markers) is shown with quality metrics as concentric rings: genome size, gene number, sequences/contigs/chromosomes with at least 5 BUSCO genes, sequence number, N50, and genome and protein BUSCO completeness and duplication. Colours are grey-scale for descriptive metrics, or a colour-blind-safe traffic light (Good/Warn/Poor) for metrics with a defensible quality direction; both ring selection and thresholds are user-configurable (Supplementary Table S3).

In the circular layout (**Figure 2**), quality metrics are shown as concentric rings, optionally from inner to outer: genome size, gene number, number of sequences/contigs/chromosomes with at least 5 BUSCO genes, sequence number, N50, and genome and protein BUSCO completeness and duplication. Colours are grey-scale for metrics considered descriptive rather than “good” or “bad” (e.g. genome size, gene number), versus a colour-blind-safe traffic light for metrics with a defensible good/bad direction (e.g. BUSCO completeness ≥95%). Good/Warn/Poor thresholds are pre-set per phylogenetic group, and can be fully customised by the user to suit any organism (**Supplementary Table S3**).

### Interactive visualisation (Shiny app)

Because no single static tree suits every comparison and users may have preference on what is displayed, nf-core/genomeqc also outputs a self-contained executable that launches an interactive Shiny application (**Figure 3**). Within the app, users can add or remove summary statistics, rescale and reorder the annotations, and adjust the plot in real time before exporting publication-ready figures as PNG or SVG. This allows the same run to be explored from several angles without re-execution, and produces the customisable figures used throughout this manuscript. It’s also possible to choose a traditional tree layout (**Figure 3 bottom**), here the same phylogeny is instead shown left-aligned and rooted, with each genome’s statistics as a row of adjacent panels rather than rings. Unlike the circular layout, all panels here use fixed colours rather than graded thresholds, but carry additional detail not shown in the circular rings: genome size is split to show GC content as a proportion of the bar, and gene number is split to show the proportion of overlapping genes; a transposable-element composition panel breaks down repeat content per genome by category (SINE, LINE, LTR, DNA transposon, etc.); and, where decontamination was run, a panel shows the proportion of each assembly flagged as foreign by FCS-GX.

**Figure 3.**
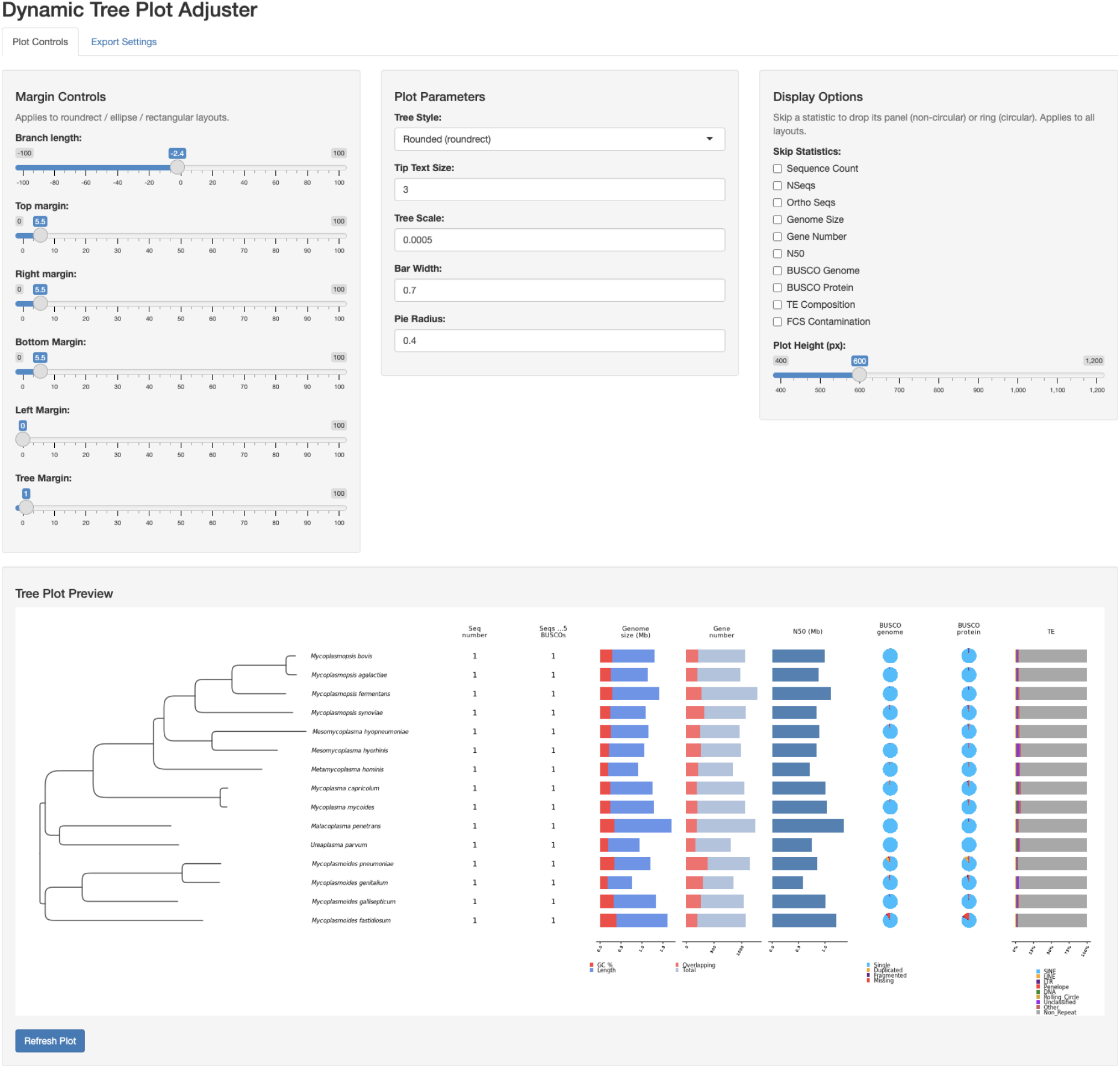
Shiny app screen shot. **TOP:** Options to adjust the figure. “Margin controls” allow the user to configure the size of the tree and margins around it. “Plot Parameters” allow a choice of tree style, the size of font, tree bar width and pie diameters. “Display options” are available to toggle off options printed onto the tree. **BOTTOM:** A rendered traditional tree. The same phylogeny as Figure 2, shown as a left-aligned rooted tree with each genome’s statistics as adjacent panels: sequence number, orthologous sequences (an estimate of chromosome number), genome size (with GC content shown as a proportion of the bar), gene number (with the proportion of overlapping genes shown as a proportion of the bar), N50, BUSCO completeness, transposable-element composition, and (where run) FCS-GX contamination.

### Additional example outputs

Alongside the comparative tree, nf-core/genomeqc produces a set of per-assembly diagnostic outputs, summarised across all species in a final HTML report (**Figure 4A**) that collates the underlying statistics from every tool that was run. Per-assembly diagnostics include a BUSCO ideogram(16) mapping the location of single-copy markers along each assembly (**Figure 4B**), in our example, the ideogram of *Mycoplasmoides pneumoniae* shows a cluster of duplicated markers near the ends of the sequences that is absent in *M. genitalium* and *M*.*fastidiosum*. The pipeline also identifies canonical telomeric repeats (where present) with tidk (27), which can distinguish assemblies that reach telomere-to-telomere contiguity from those missing terminal repeats, screens for contamination with the NCBI FCS suite(20) and classifies sequences with Tiara(21), flagging foreign, adaptor, vector and organellar sequences; and summarises transposable-element content using multiple optional tools (e.g. RepeatMasker or HITE).

**Figure 4.**
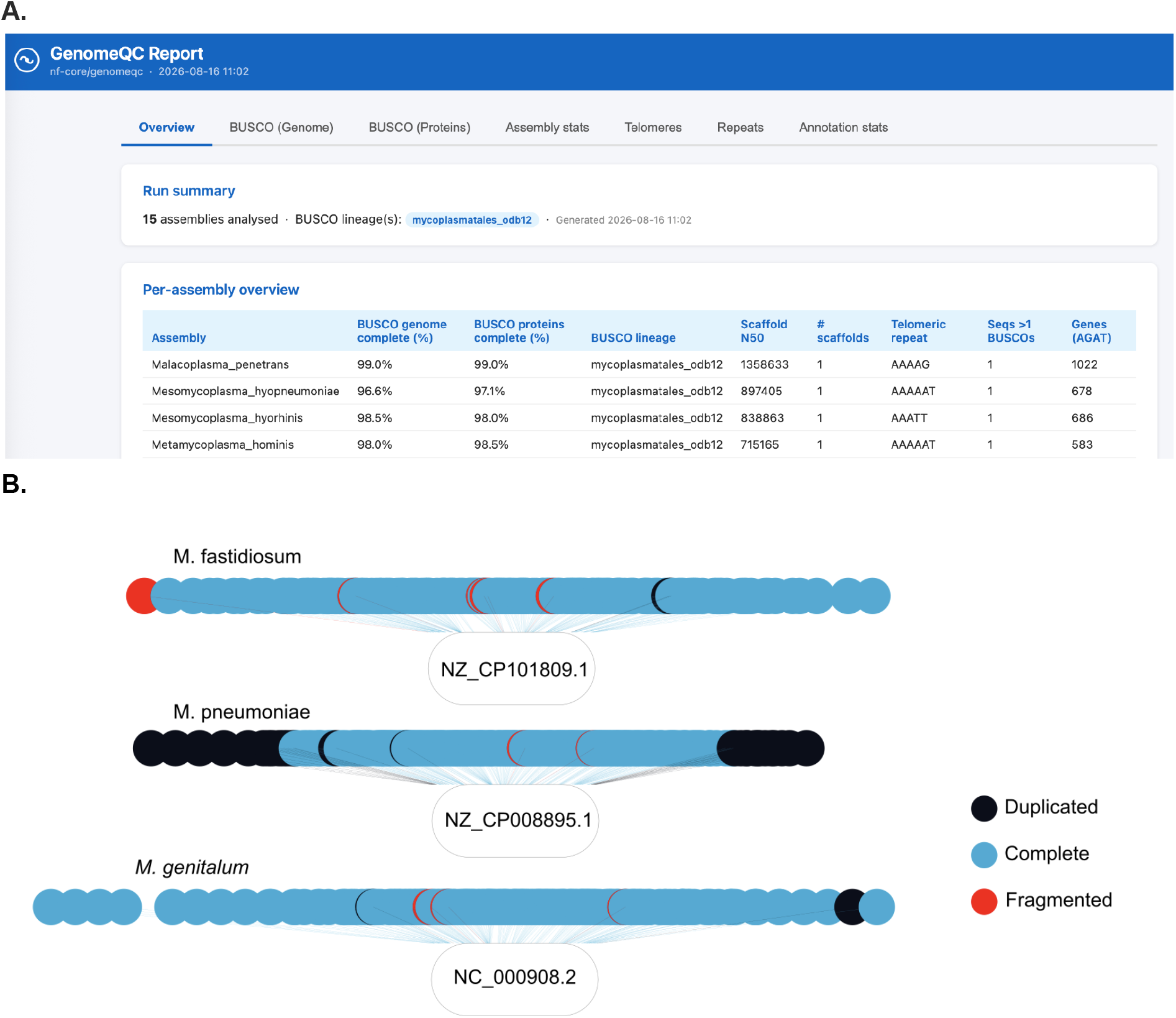
Example output. (A) Excerpt of the final HTML summary report, collating per-assembly statistics across all tools run. (B) BUSCO ideogram showing the location of single-copy markers along each assembly, shown here for three example genomes.

## Discussion

The **nf-core/genomeqc** pipeline offers a standardized solution for evaluating genome assembly and annotation quality through a single, containerized workflow. By leveraging the nf-core framework, the pipeline ensures reproducibility across computing environments while maintaining accessibility for users. Documentation and ongoing support are available through the nf-core website (https://nf-co.re/genomeqc) and via community discussion channels (https://nfcore.slack.com/channels/genomeqc) and on Github for issues (https://github.com/nf-core/genomeqc/).

A key distinguishing feature of **nf-core/genomeqc** is its unified and comparative approach to quality assessment visualization. While tools like ‘cogeqc’, ‘Blobtoolkit’, ‘AssemblyQC’, and ‘HuffordLab/GenomeQC’ provide valuable individual quality metrics and figures, nf-core/genomeqc integrates these measurements within a phylogenetic context. This design enables direct comparison of assembly statistics, gene content, telomeric features, and potential contamination across taxonomically related genomes. By organizing quality metrics along a phylogenetic tree, researchers can more readily distinguish genuine biological differences from technical limitations or assembly artifacts. The pipeline further supports flexible data input, accepting either direct NCBI database identifiers or user-provided genome files and a number of additional programs that don’t exist in current workflows.

Through its incorporation into the nf-core community framework, **nf-core/genomeqc** benefits from ongoing maintenance and will continue to evolve alongside advancing genomic technologies. Planned enhancements include custom metrics for detailing metadata (e.g. sequencer), optional choice of repetitive element annotation program, and potential integration with genome assembly workflows. The pipeline’s modular Nextflow DSL2 architecture facilitates these future developments and enables seamless incorporation of emerging quality assessment tools. By fostering collaboration within the broader genomics community, we aim to establish nf-core/genomeqc as a central resource for ensuring genomic data quality in comparative studies.

## Data availability

nf-core/genomeqc code is hosted on GitHub under the nf-core organization https://github.com/nf-core/genomeqc and released under a MIT license. The data sets used to generate the exemplary results are available within the Github repository in the input samplesheets (https://github.com/nf-core/genomeqc/assets).

## Supplementary data

None

## Acknowledgements

We thank the whole nf-core community for general support during the writing of the pipeline, and individual members who contributed to hackathons who contributed to the code base in some way including. We thank Mahesh Binzer-Panchal for early support and helpful suggestions in setting up the pipeline.

