## Supplementary Methods for "nf-core/genomeqc: a best-practice pipeline for comparing genome and assembly quality"

### Pre-processing

Before the core quality-assessment tools are run, inputs are fetched and standardised. When NCBI accessions are supplied instead of local files, genome assemblies and their annotations are downloaded with the NCBI genome download tool (assemblies from GenBank or RefSeq; annotations from RefSeq only). Annotations are parsed and standardised with the AGAT toolkit (convertspgxf2gxf) by default, or optionally with GffRead's validation mode (gffread -E, selectable via --val_tool gffread); both GTF and GFF3 input are accepted and processed by this same step.
Where annotations feed BUSCO (protein mode) completeness and OrthoFinder orthology inference, AGAT selects the longest isoform per gene, and GffRead then extracts the corresponding protein sequences in FASTA format from the assembly.

### Contamination

The pipeline can also detect contaminants from foreign organisms as well as adaptor and vector sequences using NCBI's Foreign Contamination Screen (FCS) tool suite (19). The assembly is first screened for sequences from foreign organisms, and then for exogenous adaptor and vector sequences. If contaminants are detected, affected sequences are trimmed, or removed entirely if the contaminant spans the whole sequence. This produces a cleaned assembly, but the pipeline will continue with the raw assembly unless otherwise specified in the parameters file.

Following the contamination screening, the pipeline classifies each sequence in the assembly using Tiara (28). Tiara is a deep learning tool that assigns DNA sequences to domains such as Eukaryota, Bacteria, Archaea, and identifies organelle genomes.

### Transposable elements

genomeQC supports two alternative methods for transposable element (TE) annotation, selected via --te: a fast, alignment-free method using HiTE (--te hite), or a RepeatMasker-based pipeline (--te repeatmasker), described here.
By default, a repeat library is extracted from the pipeline's default Dfam partition (v3.9) using famdb.py (RepeatMasker v4.2.2), either downloaded automatically (--RM_download_db) or supplied as a pre-staged local partition (--famdb_library). No lineage is specified by default, so the full curated Dfam family set is used; this can be restricted to a specific lineage with --famdb_lineage. De novo repeat discovery with RepeatModeler can additionally be run (--run_repeatmodeler; disabled by default, as it is computationally expensive, adding 24-48 h per genome), building a genome-specific library that is merged with the Dfam library before masking.
The resulting library is deduplicated before masking, using MMseqs2 easy-linclust (v18.8cc5c) by default (--te_clusterer; MMseqs2 easy-cluster and CD-HIT-EST are also available), at a minimum sequence identity and coverage of 0.8 by default (--te_cluster_identity, --te_cluster_coverage). When RepeatModeler is not used, deduplication runs once and the resulting library is applied identically across all genomes; when enabled, deduplication runs per genome on each genome's combined Dfam + de novo library. Each genome is then soft-masked with RepeatMasker v4.1.5, in rush mode (-qq) by default - the fastest, lowest-sensitivity setting (--repeatmasker_speed; default and quick (-q) modes are available).
Per-genome RepeatMasker .tbl summaries are parsed into eight TE categories - SINE, LINE, LTR, Penelope, DNA transposons, rolling-circle elements, unclassified, and "other" (satellites, simple repeats, low-complexity, and small RNA combined), each expressed as a percentage of total genome length, as reported directly by RepeatMasker. The non-repetitive fraction is calculated as the

complement (100% minus the summed TE percentage), rather than taken from RepeatMasker's output directly.

**Completeness**

BUSCO assesses the completeness of genome assemblies and annotations by evaluating the presence of expected universal single-copy orthologs present in their database (24). If the genome assembly is the only input, BUSCO runs on genome mode, and the genome completeness is assessed. If the pipeline runs on both genome assembly and annotation, the completeness can be assessed on BUSCO's genome mode or on protein mode, depending on the parameter selection. If protein mode is set, BUSCO will run on the extracted longest isoforms, thus evaluating annotation completeness instead.

Merqury is a reference free evaluation tool for genome assemblies. It uses k-mers from the raw unassembled reads, and compares them to those present in the genome assembly to assess their accuracy and completeness (25). It requires high-quality sequencing reads from the corresponding assembly to run. Genomeqc only works with pseudo-haplotype assemblies, and therefore this Mercury implementation does not perform phasing assessment.

Another way to assess genome completeness is through the presence or absence of telomeric repeats (26). In this pipeline, genome assemblies are screened for the presence of candidate telomeric repeats using the telomere identification toolkit (tidk), a tool that detects and reports canonical telomeric motifs in *de novo* assemblies. Once these motifs are found, summary statistics are reported, and results can be visualised.

**Circular QC plot options**

Using nf-core/genomeqc's circular tree summary (--tree_style circular), which draws sequence count, scaffold N50, BUSCO completeness, BUSCO duplication, and where decontamination was run FCS-GX contamination as concentric rings around the phylogeny, each coloured by a discrete Good/Warn/Poor score (**Supplementary Table S3**).
Because contiguity expectations vary enormously across taxa (a bacterial N50 of 1 Mb is excellent, while the same value would be poor for a vertebrate genome), scoring is controlled by a --quality_preset selecting one of six built-in threshold sets tuned to broad phylogenetic groups: generic (default, deliberately lenient), vertebrate, insect, plant, fungi, and bacteria. Thresholds are fixed reference values rather than being computed from the distribution of samples in a given run, so that a genome's grade reflects an absolute, reproducible standard and does not shift depending on which other genomes happen to be analysed alongside it in the same run.
Where a built-in preset does not match a project's expectations, individual cut-offs can be overridden without replacing the whole preset, via --quality_thresholds, a comma-separated list of metric=good:warn pairs, e.g.: --quality_preset bacteria --quality_thresholds 'n50=2e6:5e5,seq_number=50:500'. Only the numeric cut-offs can be customised; whether a metric is "higher is better" (e.g. BUSCO completeness) or "lower is better" (e.g. sequence count) is fixed per metric and cannot be altered, preventing a misconfigured threshold from silently inverting a metric's interpretation. The same custom thresholds can be set interactively through the accompanying Shiny application, via a "Custom" option in the quality-preset selector.
