## Supplementary Tables for "nf-core/genomeqc: a best-practice pipeline for comparing genome and assembly quality"

**Table S1. Comparison of tools used in this pipeline compared to other existing software/pipelines.** Showing features of existing pipelines in comparison to nf-core/genomeqc v1.0.0, which are HuffordLab/GenomeQC(1), cogeqc(2), BlobToolKit(3) and AssemblyQC(4). Only the latter two and nf-core/genomeqc have been implemented in Nextflow/Snakemake, making them equally scalable, automated and easy to use.

**Table S1:** Comparison of tools and characteristics of genome quality tools/pipelines

| **Feature** | **HuffordLab/ GenomeQC** (1) | **Cogeqc** (2) | **BlobToolKitsV0.9.0 -Scyther** (3) | **AssemblyQC V3.0.0** (4) | **nf-core/ genomeQC v.1.0.0** |
|---|---|---|---|---|---|
| **Workflow management system** | **None** | **R package** | **Snakemake/ Nextflow** | **Nextflow** | **Nextflow** |
| **Completeness (BUSCO)**(5) | **Yes** | **Yes** | **Yes** | **Yes** | **Yes** |
| **Foreign Seq Contamination**(6) | **No** | **No** | **Yes (busco/blast)** | **Yes (fsc)** | **Yes (fsc)** |
| **LTR contiguity (LAI)** | **Yes** | **No** | **No** | **Yes** | **No** |
| **Vector/Adaptor contamination** | **Yes** | **No** | **No** | **Yes** | **Yes** |
| **Telomere repeat identification (TIDK)** | **No** | **No** | **No** | **Yes** | **Yes** |

| | | | | | |
|---|---|---|---|---|---|
| Merqury (Kmer Completeness) | No | No | No | Yes | Yes |
| Transposable element classification | No | No | No | No | Yes |
| Scaffold statistics (N50/N90) | Yes (custom scripts) | Yes | Yes | Yes | Yes |
| GC content | No | Yes | Yes | Yes | Yes |
| Phylogenetics (orthofinder) | No | Yes (partial) | No | Yes | Yes |
| Phylogenetic quality summary | No | Yes (partial) | No | No | Yes |
| Metagenomics: Microbe quantificiation (kraken 2) | No | No | No | Yes | No |
| HiC QC | No | No | No | Yes | No |
| BUSCO ideograms | No | No | No | No | Yes |
| Orthologous Chromosome/ Contig number | No | No | No | No | Yes |
| Input validation | No | No | No | Yes | Yes |
| Gene overlaps | No | No | No | No | Yes |
| Genome + annotation | Yes | Yes | No | Yes | Yes |
| Gene structure statistics (e.g. AGAT) | Yes | Yes | No | Yes | Yes |
| Syntenic relationships | No | No | No | Yes (mummer) | No |
| Diploid/Haploid | No | No | No | Yes | No |
| MultiQC(7) | No | No | No | No | Yes |

**Supplementary Table S2. Comma separated samplesheet for mycoplasma species**

**input.** Showing NCBI Refseq IDs for each species, and omitting the fasta and gff (gene feature format) input (which are generated from the Refseq download) and fastq inputs which we do not have access to. “Taxid” is provided for each species, to allow fcs (foreign contamination screen) to know what is likely a contaminant in the genome.

species,ncbi,fasta,gff,fastq,taxid
Mycoplasmoides_genitalium,GCF_000027325.1,,,,243273
Mycoplasmoides_pneumoniae,GCF_000733995.1,,,,1441379
Mycoplasmoides_gallisepticum,GCF_017654545.1,,,,2096
Mycoplasmoides_fastidiosum,GCF_024498275.1,,,,92758
Mycoplasma_mycoides,GCF_018389705.1,,,,40477
Mycoplasma_capricolum,GCF_000012765.1,,,,340047
Mycoplasmopsis_bovis,GCF_000183385.1,,,,289397
Mycoplasmopsis_agalactiae,GCF_000063605.1,,,,347257
Mycoplasmopsis_synoviae,GCF_000969765.1,,,,1267001
Mesomycoplasma_hyopneumoniae,GCF_000008205.1,,,,262719
Mesomycoplasma_hyorhinis,GCF_900476065.1,,,,2100
Metamycoplasma_hominis,GCF_000767725.1,,,,1267000
Malacoplasma_penetrans,GCF_000011225.1,,,,272633
Mycoplasmopsis_fermentans,GCF_000186005.1,,,,943945
Ureaplasma_parvum,GCF_000019345.1,,,,505682

**Supplementary Table S3**. Quality-scoring thresholds for the circular tree summary

| Metric | Direction | Preset | Good | Warn | Poor |
|---|---|---|---|---|---|
| **BUSCO complete (S+D %)** | higher is better | all presets | >= 95% | 90-95% | < 90% |
| **BUSCO duplicated (%)** | lower is better | generic, vertebrate, insect, fungi, bacteria | <= 5% | 5-10% | > 10% |
| **BUSCO duplicated (%)** | lower is better | plant | <= 10% | 10-20% | > 20% |
| **Scaffold N50 (bp)** | higher is better | generic, insect, plant, fungi | >= 1 Mb | 100 kb - 1 Mb | < 100 kb |
| **Scaffold N50 (bp)** | higher is better | vertebrate | >= 10 Mb | 1-10 Mb | < 1 Mb |
| **Scaffold N50 (bp)** | higher is better | bacteria | >= 500 kb | 100-500 kb | < 100 kb |
| **Sequence count (contigs/scaffolds)** | lower is better | generic, vertebrate, insect | <= 1000 | 1,000-10,000 | > 10,000 |
| **Sequence count (contigs/scaffolds)** | lower is better | plant | <= 5000 | 5,000-50,000 | > 50,000 |
| **Sequence count (contigs/scaffolds)** | lower is better | fungi | <= 100 | 100-1000 | > 1000 |
| **Sequence count (contigs/scaffolds)** | lower is better | bacteria | <= 10 | 10-100 | > 100 |
| **FCS-GX non-contaminant (%)** | higher is better | all presets (fixed) | >= 99.5% | 98-99.5% | < 98% |

**Supplementary table 3.** In the circular tree layout (--tree_style circular), five QC metrics are scored into a discrete Good/Warn/Poor "traffic light" (colour-blind-safe Okabe–Ito palette: Good #009E73, Warn #E69F00, Poor #D55E00) and drawn as concentric rings. Each metric is compared against fixed numeric cut-offs, chosen from one of six built-in presets (--quality_preset), or overridden individually by the user (--quality_thresholds). Thresholds are not computed relative to the other genomes in a given run — they are fixed reference values, so a genome's grade is reproducible regardless of what else is analysed alongside it. BUSCO completeness cut-offs (95%/90%) follow community convention and are the same for all presets. FCS-GX contamination cut-offs are deliberately identical across every preset: there is no taxon-dependent expectation for how much of a genome should be foreign sequence, unlike assembly contiguity. N50 and sequence-count cut-offs are clade-dependent starting points (a 1 Mb N50 is unremarkable for a bacterial genome but poor for a vertebrate one) and are intended to be tuned per project rather than treated as fixed community standards. Generic (the default) is deliberately lenient, using the least strict N50/sequence-count combination, for use when no clade-specific preset applies. Descriptive metrics shown as inner rings (genome size, gene number, sequences with ≥5 BUSCOs, orthologous sequence count) are not scored — they use a neutral grey intensity ramp rather than a threshold, since they have no universal "good" direction.
